# Forward genetic screen of homeostatic antibody levels in the Collaborative Cross identifies MBD1 as a novel regulator of B cell homeostasis

**DOI:** 10.1101/2022.08.30.505889

**Authors:** Brea K. Hampton, Kenneth S. Plante, Alan C. Whitmore, Colton L. Linnertz, Emily A. Madden, Kelsey E. Noll, Samuel P. Boyson, Breantie Parotti, Timothy A. Bell, Pablo Hock, Ginger D. Shaw, Fernando Pardo-Manuel de Villena, Martin T. Ferris, Mark T. Heise

## Abstract

Variation in immune homeostasis, the state in which the immune system is maintained in the absence of stimulation, is highly variable across populations. This variation is attributed to both genetic and environmental factors. However, the identity and function of specific regulators have been difficult to identify in humans. We evaluated homeostatic antibody levels in the serum of the Collaborative Cross (CC) mouse genetic reference population. We found heritable variation in all antibody isotypes and subtypes measured. We identified 4 quantitative trait loci (QTL) associated with 3 IgG subtypes: IgG1, IgG2b, and IgG2c. While 3 of these QTL maps to known immunologically important regions of the genomes, *Qih1* (associated with variation in IgG1) mapped to a novel locus on Chromosome 18. We further associated this locus with B cell proportions in the spleen and show that Methyl-CpG binding domain protein 1 is a novel regulator of homeostatic IgG1 levels in the serum and marginal zone B cells (MZB) in the spleen, consistent with a role in MZB differentiation to antibody secreting cells.

**Summary:** Homeostatic antibody levels are highly variable between Collaborative Cross (CC) mouse strains. This forward genetic screen in the CC identifies Methyl-CpG binding domain protein 1 (MBD1) as a regulator of homeostatic IgG1 levels and marginal zone B cell differentiation.

## Introduction

Immune homeostasis is the stable state that the immune system maintains in the absence of insult. While the majority of studies on host immunity have focused on the response to specific stimuli (e.g., pathogens, vaccines, allergens, or adjuvants), a growing body of evidence suggests that an individual’s baseline immune status affects subsequent innate or antigen specific immune responses (Gnjatic et al., 2017; Graham et al., 2019; HIPC-CHI Signatures Project TeamHIPC-I Consortium, 2017; Tsang et al., 2014). Immune homeostatic parameters are gaining increasing recognition as predictors of clinical outcomes to immunotherapy and vaccination (Gnjatic et al., 2017; HIPC-CHI Signatures Project TeamHIPC-I Consortium, 2017), and several studies have shown that dysregulation of immune homeostasis can contribute to the development of cancer, autoimmunity, allergies, as well as the progression of immune-related pathology in response to infection (Crimeen-Irwin et al., 2005). Natural antibodies, which are present before antigen exposure, are a major component of immune homeostasis. These antibodies have been shown to be produced by specific subsets of B cells (e.g., marginal zone B cells and B1 B cells) and provide a first line of defense against infection through low affinity binding to pathogens (Palma J., et al. 2018). Concurrent with our understanding of B cell and antibody biology, there is growing evidence that immune homeostasis is under genetic control (Graham et al., 2017; Krištić et al., 2018; Phillippi et al., 2013; Collin et al., 2019). However, we still do not fully understand the role that genetic differences between individuals play in driving baseline immunity. Many studies have shown that humans exhibit significant inter-individual variation in baseline immune phenotypes (Cassidy et al., 1974; Grundbacher, 1974; HIPC-CHI Signatures Project TeamHIPC-I Consortium, 2017; Tsang et al., 2014). However, it has been difficult to identify the genes that contribute to this variation due to a variety of confounding factors. These include difficult to control environmental factors, such as prior microbial and environmental exposures as well as age dependent changes in immune status that create a highly dynamic immune environment (Poon M., et al. 2021). Rodent models, which allow for greater control of host genetics and environmental exposures, represent an attractive system with which to model and investigate those factors driving the development of various immune homeostatic states. Gene specific knockout mice have been critical to our understanding of the immune system. However, these models represent extreme genetic perturbations, rather than the more subtle effects on gene expression or function more commonly associated with naturally occurring genetic variation in humans.

To better model how natural genetic variation impacts immune homeostasis, we turned to the Collaborative Cross (CC) genetic reference panel (grp; Collaborative Cross Consortium, 2012). The CC grp is a large set of recombinant inbred (RI) mouse strains derived from 8 founders: five classical laboratory strains (C57BL/6J (B6), A/J, 129S1/SvImJ (129S1), NOD/ShiLtJ (NOD), and NZO/HlLtJ (NZO)) and three wild-derived strains (PWK/PhJ (PWK), CAST/EiJ (CAST), and WSB/EiJ (WSB)) (Collaborative Cross Consortium, 2012). These eight founder strains capture >90% of common genetic variation present in laboratory mouse strains and represent the three major subspecies of *Mus musculus* (Threadgill et al., 2011; Welsh et al., 2012; Roberts et al., 2007). Importantly, this genetic variation exists across the genome, ensuring no blind spots for genetic mapping analyses. This genetic diversity has resulted in discovery of several genetic loci associated with a variety of biomedically relevant traits (Ferris et al., 2013, He et al., 2021; Graham et al., 2021; Gu et al., 2020; Smith et al., 2019; Noll et al., 2020; Hampton et al., 2021, *Preprint*). We and others have shown that there is extensive variation in splenic T cell populations (Graham et al., 2017), antibody glycosylation patterns (Krištić et al., 2018), as well as variation in the general immune landscape of the spleen (Collin et al., 2019), across the CC population.

Here, we investigated how genetic variation impacts one aspect of systemic immune homeostasis: baseline serum antibody levels. We found that serum antibody levels for a variety of immunoglobulin (Ig) isotypes was highly varied across the CC, and we identified four quantitative trait loci (QTL) regulating these phenotypes. Three of the identified QTL mapped to known immunologically relevant regions of the genome, including the major histocompatibility locus and the immunoglobulin heavy chain locus. We also identified a novel locus broadly associated with variation in serum antibody levels, as well as B cell subset proportions in the spleen. Further analysis and independent mutant generation showed that the Methyl-CpG binding domain protein 1 (*Mbd1*) regulates homeostatic antibody levels and B cell differentiation, providing new insights into the genetic regulation of the humoral immune system and highlighting the utility of forward genetics-based discovery approaches in studying immunity.

## Results

### Variation in baseline antibody levels in the Collaborative Cross (CC) is largely driven by genetic factors

We quantified antibody concentrations in the serum of 117 mice from 58 CC strains (median = 3 mice/strain, 6-8 weeks old) to understand the role of genetic variation on homeostatic antibody levels. Serum from these mice was collected seven days after a non-specific footpad infection of phosphate buffered saline (PBS) with 1% fetal bovine serum (FBS), as these animals were control animals for another study. We assessed the concentrations of total IgA, IgM, and IgG, as well as IgG subtypes (IgG1, IgG2a, IgG2b, IgG2c, IgG3) in these sera by ELISA. We found that antibody concentrations varied greatly (3-50-fold) across these mice (**Figure 1**). For example, IgM levels varied from 7.86 ug/mL – 271.39 ug/mL, total IgG levels ranged from 58.76 ug/mL – 173.89 mg/mL, and IgA varied from 4.39 ug/mL – 257.9 ug/mL in this population. Importantly, the variation between strains is much greater than that within strains (median standard deviation within strains is 0.596 ug/mL across all isotypes). We more formally assessed this by estimating the broad sense heritability (proportion of phenotypic variance in our population which can be attributable to genetic differences between individuals) for each of our antibody phenotypes and found that heritability estimates ranged from 0.25 – 0.66 (**Table 1**), indicating that genetic differences between strains plays a strong role in impacting this observed phenotypic variation in the CC. To understand the phenotypic relationships between antibody isotypes at homeostasis, we looked at pairwise correlations between all antibody isotypes and subtypes measured in the serum. We found that several antibody isotypes were highly correlated (mean correlation coefficient: 0.406, range: -0.115 – 0.841). We find that IgG1 levels were highly correlated with total IgG levels (correlation coefficient = 0.805), which was expected given the well characterized prevalence of IgG1 in the makeup of total IgG. However, we found additional interesting relationships among other antibody isotypes. For example, we found that IgA was highly correlated with both IgG2a (correlation coefficient = 0.645) and IgG2b (correlation coefficient = 0.543) (**Figure 1**). These data suggests that there may be common genetic regulation of homeostatic antibody levels.

**Table 1:** Homeostatic antibody serum concentration ranges and broad sense heritability estimates.

| Antibody isotype | Phenotypic range (ug/mL) | Median Phenotype Value (ug/mL) | Heritability Estimate |
| --- | --- | --- | --- |
| Total IgG | 58.76 – 173895.19 | 1498.06 | 0.488 |
| IgG1 | 78.09 – 280910.17 | 1834.49 | 0.518 |
| IgG2a | 10.71 – 2378.67 | 187.10 | 0.462 |
| IgG2b | 3.07 – 5448.50 | 149.16 | 0.641 |
| IgG2c | 0.29 – 536.33 | 79.45 | 0.661 |
| IgG3 | 5.61 – 2906.31 | 101.02 | 0.249 |
| IgM | 7.87 – 271.39 | 44.22 | 0.561 |
| IgA | 4.39 – 257.93 | 41.63 | 0.375 |

**Figure 1:**
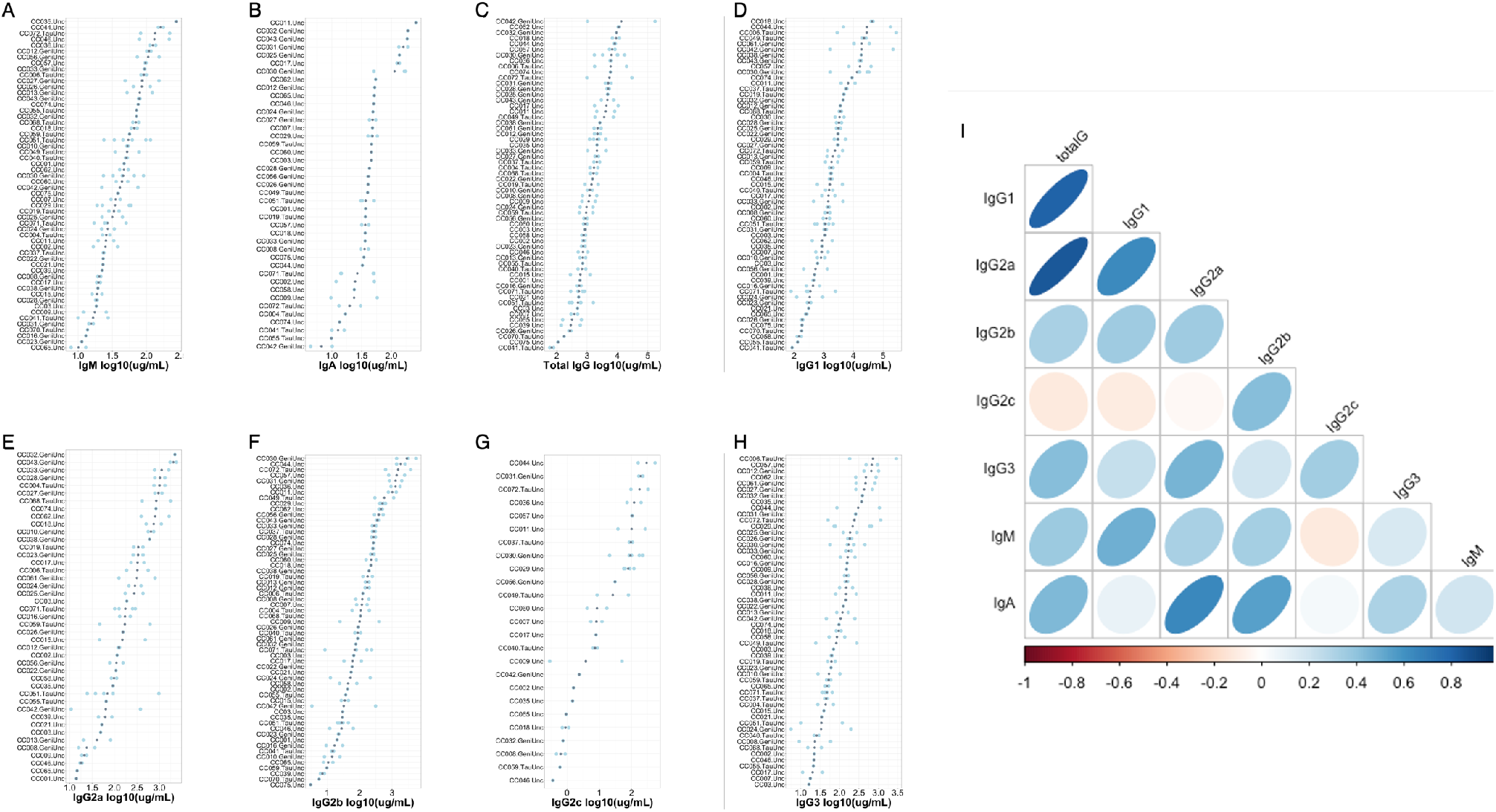
Variation in baseline antibody levels across CC strains and antibody isotypes. Total antibody levels for each of IgG subtypes, total IgG, IgA, and IgM were quantified by ELISAs. (A-H) Each plot is individually ordered so that the strain with the greatest strain mean is at the top of the plot and the strain with the lowest is at the bottom. Grey dots indicate the mean antibody concentration for an individual strain, and light blue dots indicate individual animal measurements. (I) the relationship between individual antibody isotypes and subtypes in the CC strains.

We next conducted genetic mapping to identify polymorphic genome regions associated with antibody level differences between the strains. This quantitative trait locus (QTL) mapping identified four genome regions associated with variation in antibody levels (**Table 2**). *Qih1*, QTL for immune homeostasis 1 (Chromosome 18:73 – 78Mb, genome-wide p < 0.2), was identified for variation in homeostatic IgG1 levels, with the B6, CAST, and WSB haplotypes associated with higher levels of homeostatic IgG1. *Qih2* (Chromosome 17: 43.46 – 44.03Mb, genome-wide p < 0.05) was identified for variation in homeostatic IgG2c levels, where NOD, B6, CAST, and NZO haplotypes were all associated with higher levels of IgG2c. Additionally for IgG2c, we identified *Qih3* (Chromosome. 12:117.73 – 120.02Mb, genome-wide p < 0.2), driven by the B6 and NOD haplotypes, which are associated with higher levels of IgG2c. Lastly, we found *Qih4* (Chromosome 12: 112.93 – 115.06Mb, genome-wide p < 0.05) for variation in IgG2b levels at homeostasis. At this locus, B6 and NOD haplotypes are associated with higher levels of IgG2b. CAST, WSB, A/J, 129S1, and NOD haplotypes are intermediate, and the PWK haplotype is associated with lower levels of IgG2b.

**Table 2:**
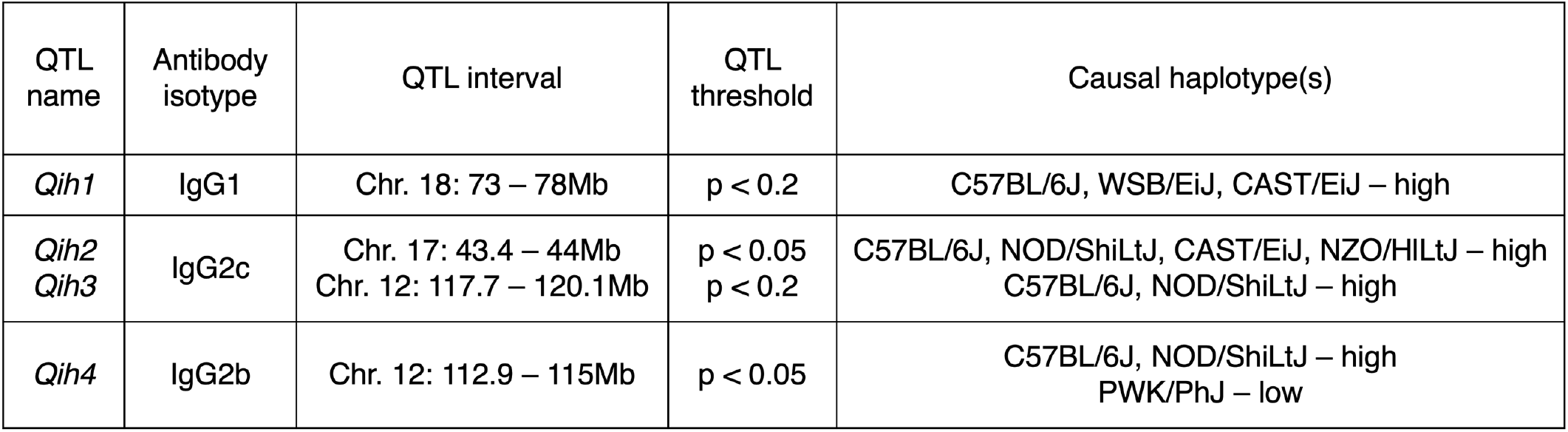
QTL identified in homeostatic antibody screen and causal haplotypes driving each QTL.

| QTL name | Antibody isotype | QTL interval | QTL threshold | Causal haplotype(s) |
| --- | --- | --- | --- | --- |
| *Qih1* | IgG1 | Chr. 18: 73 – 78Mb | $p < 0.2$ | C57BL/6J, WSB/EiJ, CAST/EiJ – high |
| *Qih2*<br>*Qih3* | IgG2c | Chr. 17: 43.4 – 44Mb<br>Chr. 12: 117.7 – 120.1Mb | $p < 0.05$<br>$p < 0.2$ | C57BL/6J, NOD/ShiLtJ, CAST/EiJ, NZO/HILtJ – high<br>C57BL/6J, NOD/ShiLtJ – high |
| *Qih4* | IgG2b | Chr. 12: 112.9 – 115Mb | $p < 0.05$ | C57BL/6J, NOD/ShiLtJ – high<br>PWK/PhJ – low |

It has long been suggested that expression of IgG2a and IgG2c is mutually exclusive in mouse strains based on the expression of IgG2a in BALB/c mice and IgG2c in B6 and SJL mice (Morgado et al., 1989; Zhang et al., 2011). However, whether the genes controlling these isotypes were physically linked on the same chromosome, or unique alleles at a locus was not understood. Recent work using isotype specific PCR showed that IgG2a and IgG2c were mutually exclusive in 4 inbred strains of mice: C57BL/6, NMRI, DBA2, and SJL, which the authors took as strong evidence of allelism (Zhang et al. 2011). Given that we mapped a QTL for IgG2c levels to the heavy chain locus (where the putative allelic variants of IgG2a and IgG2c exist), we examined these phenotypes and haplotypes much more closely and assessed the relationship between the CC founder haplotypes at this locus, and their ability to create either IgG2a or IgG2c (**Figure 2**). Phenotypically, we found that CC strains with either the B6 or NOD haplotype at the IgH locus expressed IgG2c, while strains with the other founder haplotypes at the locus expressed IgG2a. To confirm this relationship more formally with previous analyses in the literature, we queried whole genome sequences of 24 CC strains (Srivastava 2017, Shorter 2018) (3 strains with each of the founder haplotypes at IgH) using probes that would specifically identify the previously proposed IgG2c or IgG2a alleles (Morgado et al. 1989). We found that strains with B6 and NOD haplotypes had no evidence for the IgG2a probe and only had genome sequences for the IgG2c allele. Similarly, strains with the 129S1, A/J, NZO, CAST, WSB, or PWK haplotypes at IgH only had genome sequences for the IgG2a allele. Under *Qih3*, which sits at the IgH locus, the B6 and NOD haplotypes were associated with high levels of IgG2c (while also showing no evidence for IgG2a), while the other 6 founder haplotypes were associated with low/nonexistent IgG2c, as well as high IgG2a levels. We have 3 CC strains in our screen with informative recombination in this region, CC017 transitions from A/J to NOD at 118Mb, CC058 was segregating B6 and CAST alleles across the entire region, and CC060 transitions from B6 to CAST at 117Mb. We found that CC017 and CC058 animals express only IgG2a, while CC060 animals express IgG2c. As such, our data strongly support the hypothesis that IgG2a and IgG2c represent products of two alleles of the same gene, distal to 118 Mb (consistent with the genomic location of the IgG2a/c gene). Concurrently, our ability to genetically map and associate these phenotypes to a prior implicated causal locus validates our larger genetic mapping approach.

**Figure 2:**
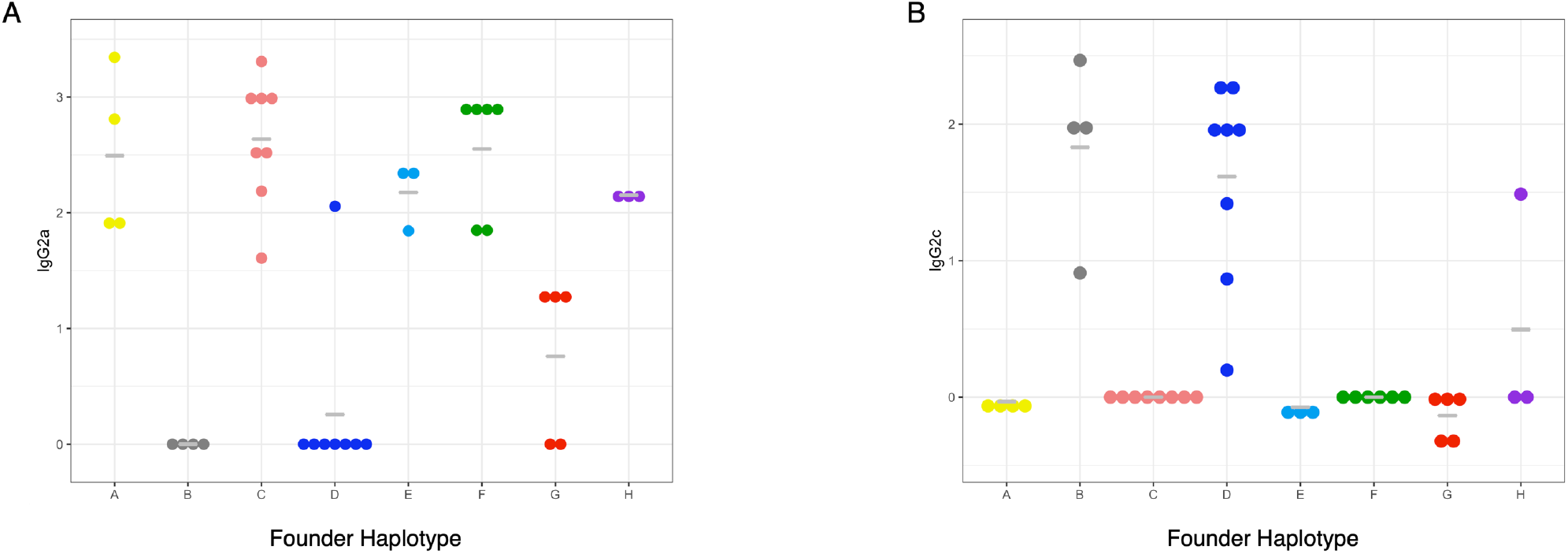
Expression of IgG2a and IgG2c by founder strain haplotype at the heavy chain locus. Grey bar indicate haplotype means and each individual point represents the strain mean for each CC strain with a given haplotype at the heavy chain locus (Chromosome 12: 113 – 120Mb). The y-axis is the log10 transformed concentration of the specified antibody isotype.

### *Qih1* broadly impacts homeostatic antibody and splenic B cells

Given the proximity of *Qih2, Qih3*, and *Qih4* to known immunologically relevant genome regions *(Qih2* with the major histocompatibility complex (MHC), and *Qih3* and *4* with the immunoglobulin heavy chain (IgH) locus), we focused our attention on the novel *Qih1* (**Figure 3**). We first asked whether *Qih1* specifically regulated IgG1 levels (the initial trait for which we mapped the locus), or if it was more broadly associated with differences in the levels of other antibody classes and isotypes. We simplified the eight haplotype groups present in the CC into high- and low-response haplotypes at *Qih1* (BFH=high, ACDEG=low) using our previously established approach (Noll et al.). We found that there was a significant association between *Qih1* haplotype groups and total IgG (p = 1.627e-6) and IgG2b (p = 0.0024) levels, as well as marginal associations with IgG2a (p = 0.015), IgG3 (p = 0.057), and IgM (p = 0.068) levels, suggesting that *Qih1* has broad effects on antibody levels at homeostasis (**Figure 4**).

**Figure 3:**
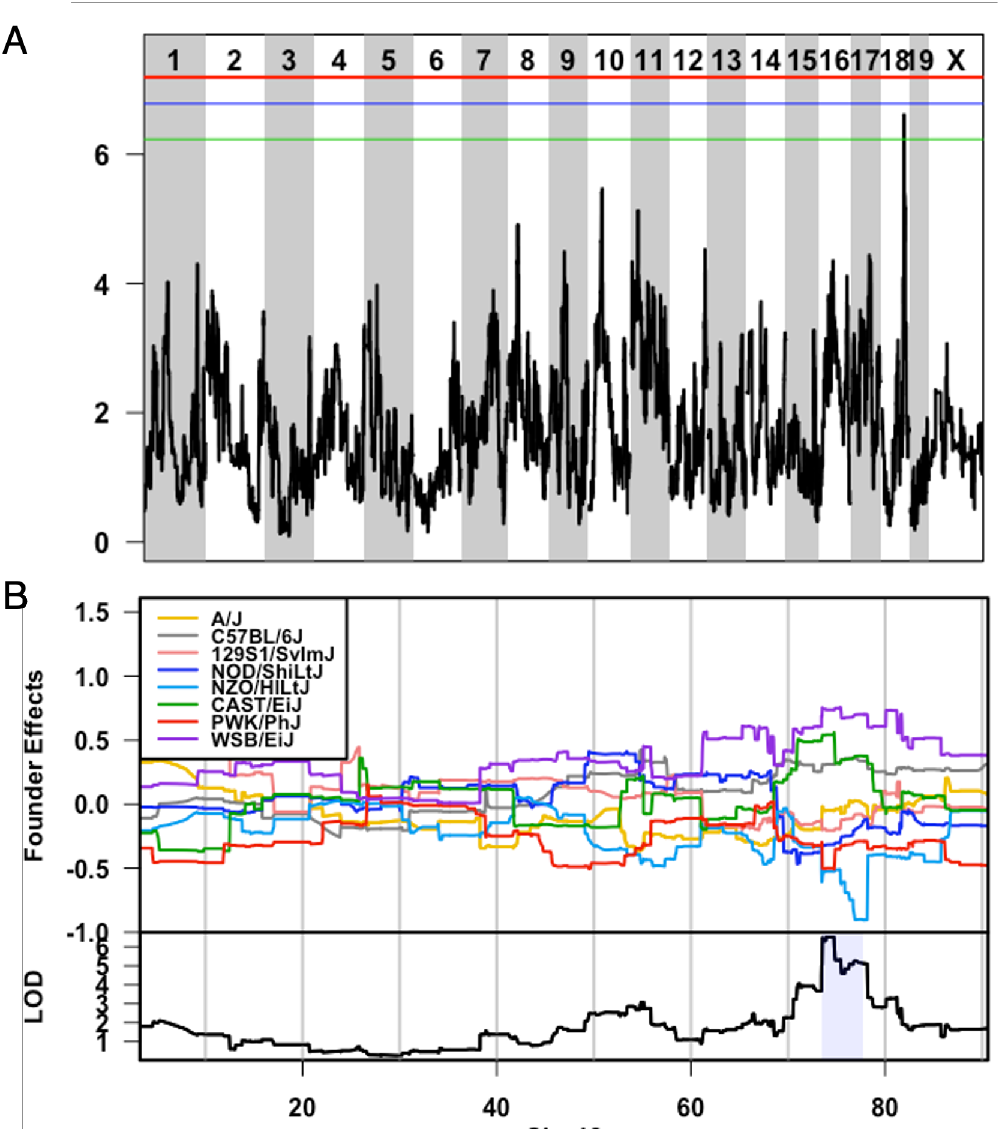
*Qih1* identified for variation in homeostatic IgG1 levels is driven by B6, WSB, and CAST haplotypes. A) LOD plot showing QTL significance (y-axis) across the genome (x-axis) with significance thresholds (genome-wide p-value = 0.05 (red), 0.1 (blue), 0.2 (green)). *Qih1* associated allele effects (A/J = yellow, B6 allele = grey, 129S1 allele = pink; NOD allele = dark blue; NZO allele = light blue; CAST = green; PWK = red; WSB = purple) were determined for the associated peak. B) Allele effect plot shows the mean deviation from population-wide mean as on the upper Y-axis for each allele segregating in the CC across the QTL peak region (x-axis positions are megabases on the chromosome) Highlighted region is the QTL confidence interval.

**Figure 4:**
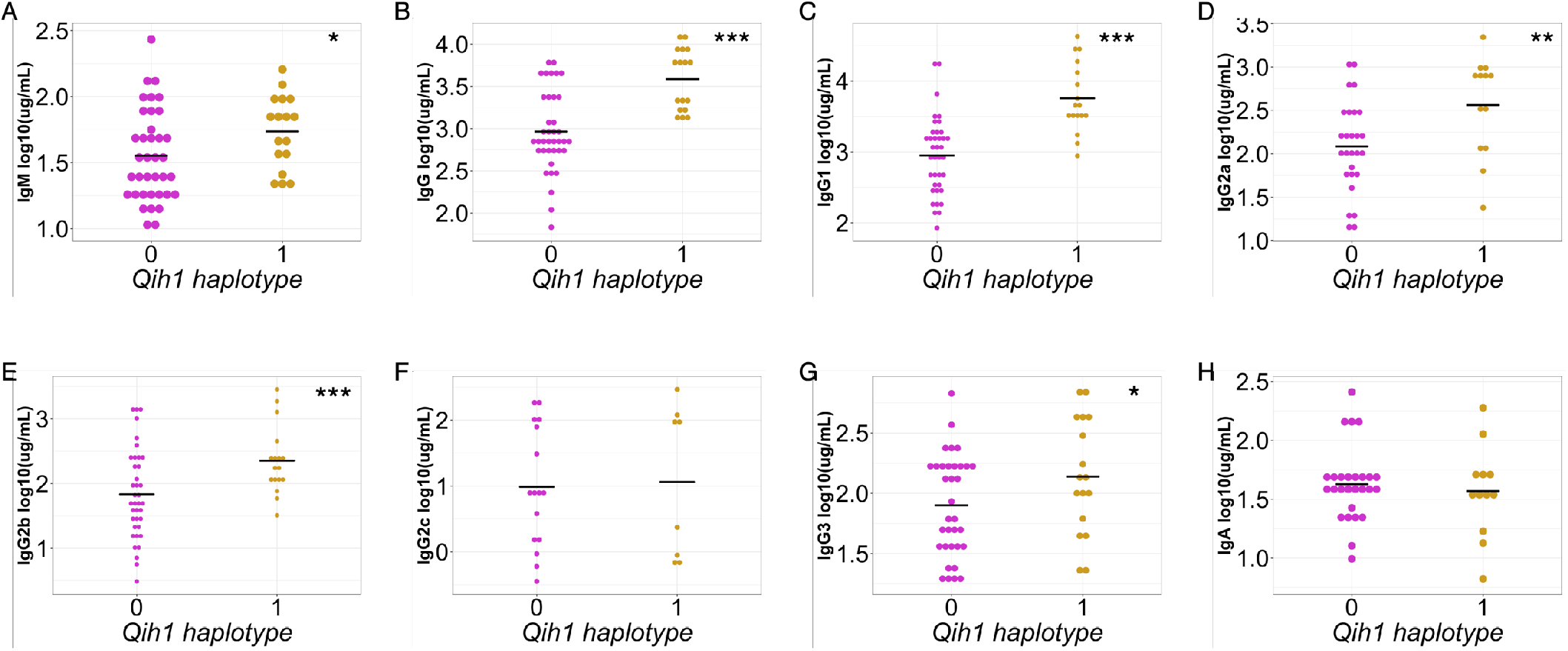
*Qih1* shows broad effects across antibody isotypes and subtypes. We assessed the relationship between B6, WSB, and CAST haplotypes (*Qih1* haplotype = 1) and other antibody isotypes and subtypes measured in our screen. Each point represents the mean value for each CC strain and the mean for each haplotype group on the x-axis is denoted by the grey crossbar. (*p < 0.1, **p < 0.05, ***p < 0.01) p-values determined using nest linear model approach described in the methods.

We next addressed whether this genetic regulation of differences in antibody levels was due to intrinsic production differences of antibody on a per cell basis or could be due to the locus controlling the abundance of specific cell populations (e.g., B cells). We took advantage of an independent cohort of 89 mice from 48 CC strains (1-2 mice per strain, supplemental table 1, Keele et al., 2020). We analyzed splenocytes from these mice for high-level immune populations (e.g., T cells, B cells, dendritic cells, and macrophages) at homeostasis, as these mice had not had any specific immunological perturbations performed on them. As with our original antibody screen, we found that these cell populations were highly variable across animals (2-6-fold differences, supplemental table 2), and that most of this variation could be attributed to differences between genotypes (heritability of 0.24-0.76, supplemental table 2). As above, we again assigned these CC strains to either the high or low haplotype groups at *Qih1* and assessed the strength of relationships between these *Qih1* haplotypes and the measured cell populations. We found that there was a significant relationship between *Qih1* and total CD19^+^ B cells in the spleen (p = 0.041), where CC strains with a high antibody level at Qih1 showed a decrease in the proportion of total splenic B cells. We also found marginal associations between *Qih1* and CD3^+^ T cells (p = 0.088) and CD8^+^ T cells (p = 0.078) in the same direction as the B cell relationship. However, we found no associations with CD4^+^ T cells (p = 0.414), CD11b^+^ cells (p = 0.169), or CD11c^+^ cells (p = 0.612) in this study (**Figure 5**). These results suggest that *Qih1* may broadly regulate multiple aspects of systemic immune homeostasis. However, consistent with its effects on antibody levels, the strongest relationship we observed was between *Qih1* haplotype and splenic B cell proportions.

**Figure 5:**
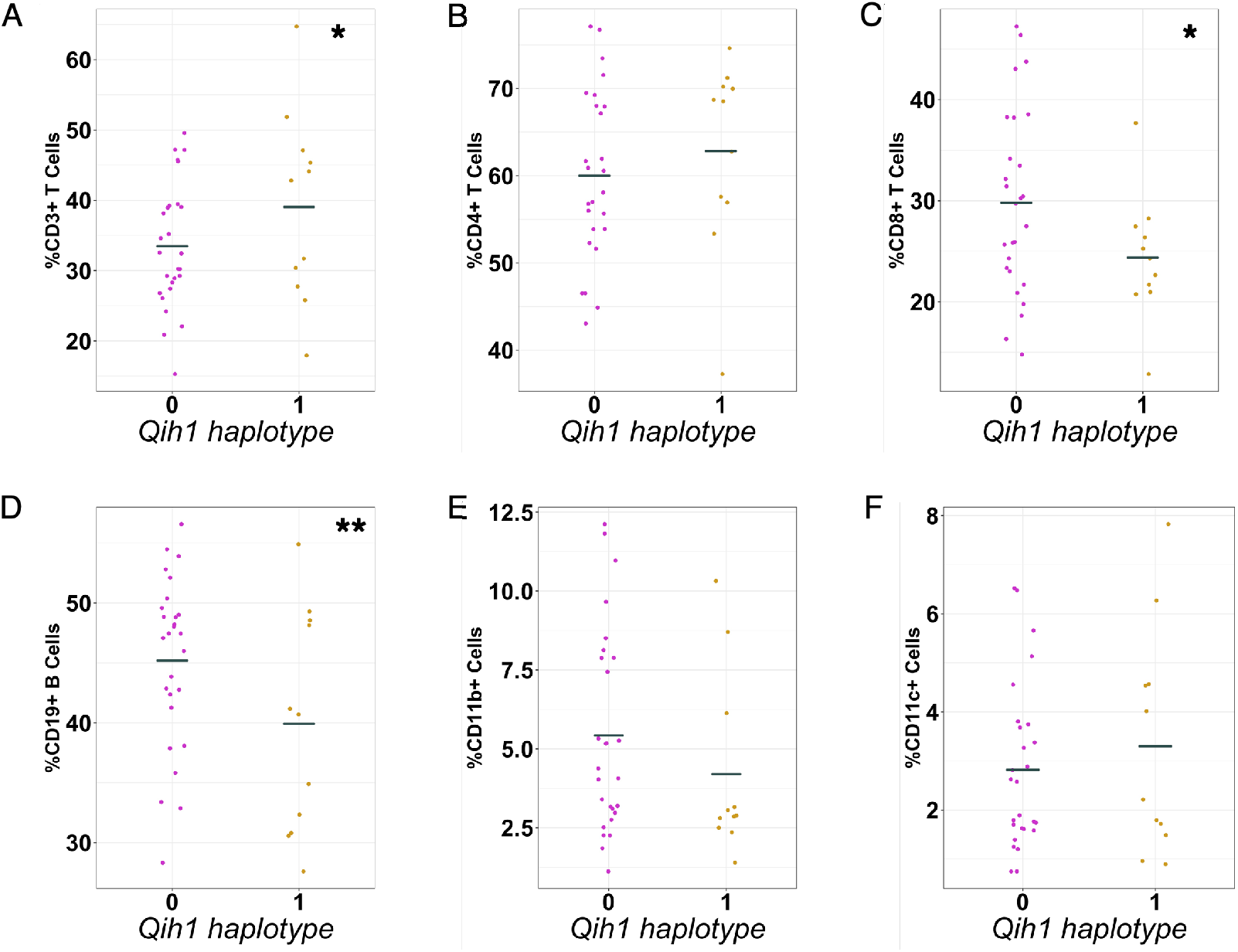
*Qih1* shows broad effects on immune cell populations in the spleen. We assessed the relationship between B6, WSB, and CAST haplotypes (*Qih1* haplotype = 1) and high-level immune cell populations in the spleen. Each point represents the mean value for each CC strain and the mean for each haplotype group on the x-axis is denoted by the grey crossbar. (*p < 0.1, **p < 0.05, ***p < 0.01) p-values determined using nest linear model approach described in the methods.

Given the relationship between *Qih1* and both antibody levels and the relative abundance of total B cells within the spleen, we next assessed how *Qih1* impacts B cell development. We assessed total serum antibody levels and several B cell subsets in the spleen and bone marrow in 67 animals from a selected a set of 12 CC strains (2-8 mice/strain, 6 strains with a B6 haplotype, and 6 strains with contrasting NZO, NOD or PWK haplotypes at *Qih1*; **supplemental table 1)**. We analyzed total B220^+^ B cells, early, immature, and mature B cells in the bone marrow, as well as total B220^+^, transitional, mature, and B1a (CD5^+^) B cells in the spleen. Across these B cell populations, we found that there was a significant relationship between splenic mature B cells and this locus (p = 0.102, **supplemental figure 1**). Concurrently, we validated the effects of this locus on total IgG and IgG1 concentrations in the serum. Taken together, these data suggest that *Qih1* may be regulating antibody levels in a B cell intrinsic manner.

### MBD1 as a novel regulator of homeostatic antibody levels and splenic B cell subsets

Concurrent with our work investigating the immune mechanisms that *Qih1* causes variation in, we used our established QTL candidate analysis pipeline (Noll et al. 2020) to identify candidate genes underlying the effects of *Qih1*. Within the *Qih1* locus, there are only 14 protein coding genes, and 10 of these genes had variants specific to all 3 causal haplotypes (in total 12 genes with CAST-specific variants, 11 genes with WSB-specific variants, and 11 genes with B6-specific genetic variants). Only one gene contained missense or nonsense (protein sequence) variants across all three haplotypes (six genes contained CAST-specific protein effecting variants, WSB and B6 specific protein effecting variants only occurred in this one gene). As such, this one gene, Methyl-CpG binding domain protein 1 (*Mbd1*) became our gene of focus.

There were not common missense variants in *Mbd1* segregating B6, CAST and WSB from the other founder strains. However, all three strains possessed independent missense variants (WSB: rs36715598; CAST: rs36834535 and rs222802617; and B6: rs46321411, rs46176119, and rs30250376). Of note was the excess of non-synonymous differences between B6 and the other common laboratory strain founders of the CC. Therefore, we further investigated the amino acid sequence variation among 29 additional mouse strains and 3 additional species (rats, non-human primates, and humans) (supplemental table 1, Keene et al., 2013; Doran et al., 2016) Besides the other Clarence Little strains (C57BL/6NJ, C57BL/10J, C57BR/cdJ, C57L/J, C58/J), only ST/bJ, BUB/BnJ (2 variants) and the wild derived ZALENDE/EiJ strains had these ‘B6’ variants. Given the otherwise high level of amino acid conservation in MBD1 across mouse strains, rats, non-human primates, and humans, this indicates that the B6 allele of *Mbd1* is both highly evolutionarily derived in this region and likely of a single wild origin that was only introduced into a subset of mouse inbred strains.

MBD1 has been previously shown to be involved in T cell development (consistent with our above observation that there were T cell differences in some of our CC analyses) and autoimmunity (Waterfield et al., 2014), neural development (Lax et al., 2017; Jobe et al., 2017), and adipocyte differentiation (Matsumura et al., 2015). While *Mbd1* deficient animals have been previously generated, these mice were initially generated on the 129S4 genetic background (Zhao X et al., 2003), and then backcrossed to C57BL/6. Given that 129 (129s1 in the case of the CC) and C57BL/6 had opposing haplotype effects, this makes it difficult to differentiate whether effects on baseline antibody or B cell populations are due to the lack of *Mbd1*, or other variants in the locus. Therefore, we generated a new *Mbd1* knockout (KO) directly on the C57BL/6J genetic background by CRISPR gene editing to directly test whether MBD1 specifically plays a role in regulating homeostatic antibody levels and more broadly, B cell subset differences.

Using a heterozygote-x-heterozygote breeding design, we found that early adult (6-8wk old) *Mbd1* KO animals had several antibody and B cell related differences relative to their WT littermates. Specifically, mutant mice had lower levels of IgG1 in the serum compared to WT littermates (Figure 6, recapitulating our observations in the CC). They also had significant increases in marginal zone B cell numbers and proportions (of all B cells) in the spleen, with no differences in total, transitional, or follicular B cells (Figure 7). Given that marginal zone B cells can rapidly differentiate into antibody secreting cells and the inverse relationship between MZBs and antibody levels in our KO mouse, we measured CD138+ antibody secreting cells in the spleen. As expected, we found a decrease in antibody secreting cell abundances in the spleens of *Mbd1* KO animals, corresponding to the decrease in antibody and increase in MZBs (Figure 7).

**Figure 6:**
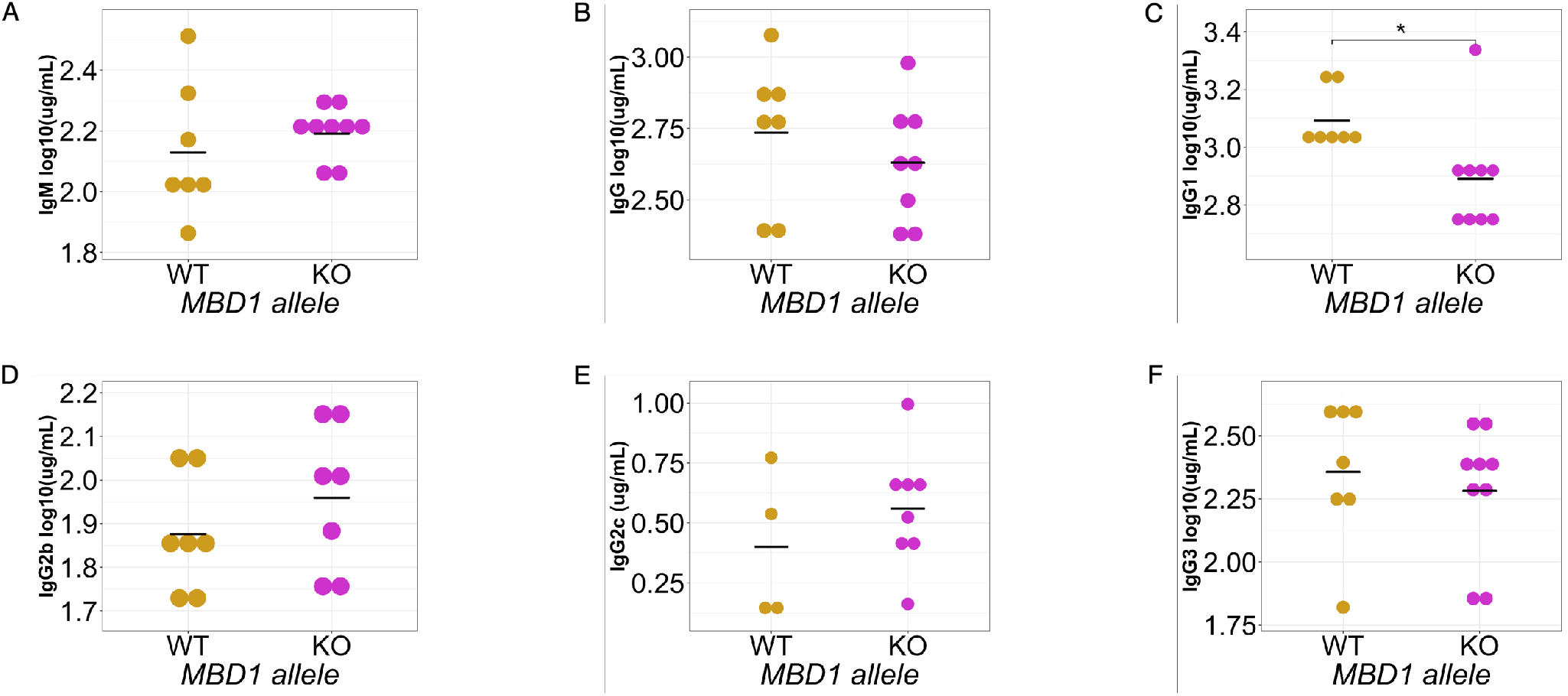
MBD1 regulates homeostatic serum IgG1 levels. The data shown represent one of two independent experiments, each performed with 6-9 KO and 6-7 WT animals. Each point represents an individual animal in the experiment and the mean for each genotype is denoted by the grey crossbar. (*p < 0.05, **p < 0.01) p-values determined using students t test or wilcox test.

**Figure 7:**
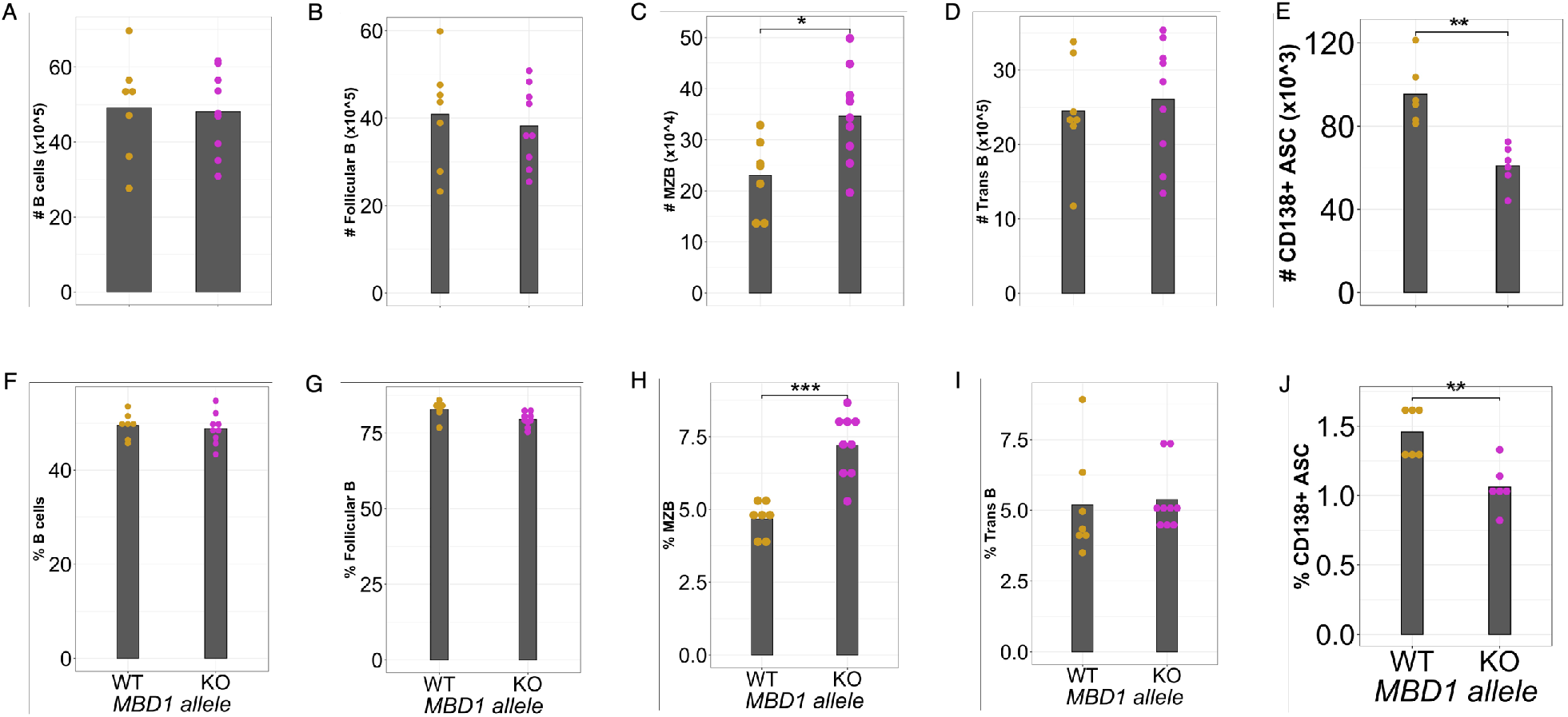
MBD1 regulates marginal zone B cell and antibody secreting cell levels at homeostasis. The data shown represent one of two independent experiments, each performed with 6-9 KO and 6-7 WT animals. Each point represents an individual animal in the experiment and the mean for each genotype is denoted by the grey bar. (*p < 0.05, **p < 0.01, ***p < 0.001) p-values determined using students t test or wilcox test.

In a separate cohort of animals, we aged them further to maturity (15-16 weeks old), and assessed IgG1, total IgG and IgM levels. We found that *Mbd1* KO animals maintained reduced levels of antibodies (total IgG and IgM, supplemental figure 2) relative to their wild type littermates through this age. All told, our data in this *Mbd1* KO stock are consistent with the haplotype effects on baseline antibody and B cell populations observed in the CC, indicate that MBD1 is a negative regulator of marginal zone B cell differentiation and a suppressor of antibody secretion. Thus, it seems likely that MBD1 acts to regulate marginal zone B cell differentiation into an antibody secreting cell, and genetic polymorphisms in the *Mbd1* gene contributing to variation in baseline antibody levels and B cell populations are due to MBD1’s role in this differentiation process.

## Discussion

Immune homeostasis is the state in which the immune system is maintained in the absence of specific insults. This balanced state is critical in ensuring an efficient response when challenged, while concurrently limiting immune pathology or unwanted (e.g. allergic) responses. The immune homeostatic state varies greatly in humans (Cassidy et al., 1974; Grundbacher, 1974; HIPC-CHI Signatures Project TeamHIPC-I Consortium, 2017; Tsang et al., 2014) and has been shown to influence responses to immunotherapies and vaccination (Gnjatic et al., 2017; HIPC-CHI Signatures Project TeamHIPC-I Consortium, 2017). Therefore, understanding the genetic mechanisms that regulate the homeostatic state is critical to understanding an individual’s potential to respond to pathogen challenge. However, it is challenging to study immune homeostasis in human populations due to environmental factors like prior pathogen exposure, diet (Kapellos et al., 2019), or environmental insults (Smeester et al., 2017) – all of which concurrently can perturb the immune system, As such, mouse models have been critical for identifying and understanding genes that regulate immune development and homeostasis (Falk et al., 1996; Kitamura et al., 1991; Lansford et al., 1998; Khattri et al., 2001; Kasprowicz et al., 2003). We extend this body work by describing variation in homeostatic antibody levels across the Collaborative Cross population and identify four QTL that are associated with variation in three different IgG subtypes (IgG1, IgG2b, IgG2c). Three of these QTL map to known immunologically important regions, *Qih2* (mapped for IgG2c) is located near the major histocompatibility locus, and *Qih3* (mapped for IgG2c) and *Qih4* (mapped for IgG2b) are located at or near the immunoglobulin heavy chain locus. We also identified a novel locus, *Qih1*, associated with variation in IgG1 levels at homeostasis, and demonstrated that this locus had broader effects on total antibody levels and splenic immune cell populations. Lastly, we identified a gene underneath *Qih1, Mbd1*, as a novel regulator of homeostatic antibody and marginal zone B cell differentiation to antibody secreting cells.

Our initial study focused on a variety of circulating antibody isotypes, and we identified loci at or near the major histocompatibility (MHC) locus and the immunoglobulin heavy chain (IgH) locus that are associated with variation in IgG2a/c or IgG2b. These loci validate our approach, as they serve as strong positive controls for regions known to be important for antibody levels. However, they can also inform potential mechanisms driving differences in total homeostatic antibody levels. *Qih4* (mapped for IgG2b), located near the IgH locus, would potentially suggest cis-regulatory elements which control expression or regulation of IgG2b alleles. *Qih2* (mapped for IgG2c), located near MHC, may indicate a haplotype specific manner of antigen detection and presentation that influences IgG2c expression or relate to the regulation of adaptive immune cell crosstalk, more generally. However, defining the specific mechanism by which *Qih2* regulates IgG2c levels requires additional analysis of T cell responses and potentially other aspects of innate immune crosstalk with the adaptive B cell compartment. More broadly, various antibody isotypes and subtypes are important for different aspects of the immune response (Collins A, 2016; Vidarsson G et al., 2014). In our study, we have identified divergent haplotypes across loci regulating antibody levels at homeostasis, which suggests that independent genetic regulation arose from antigenic exposure histories in the evolution of various mouse strains (Smith et al., 2016).

Our mapping allowed us to return to an observation long known in the literature: that laboratory mice tend to produce either IgG2a or IgG2c, but not both (Morgado et al., 1989). Additionally, even in outbred mice it was somewhat unclear if these represented two alleles or paralogues closely linked on the same chromosome (Morgado et al., 1989; Zhang et al., 2011). As described above, we found that CC strains with either B6 or NOD haplotypes at the IgH locus expressed IgG2c and strains with the other founder haplotypes at the locus expressed IgG2a. Concurrently, we took advantage of whole genome sequence data of the CC strains. We identified probe sequences from the original characterization of the IgG2a and IgG2c paralogues and confirmed for strains with a B6 or NOD haplotype at IgH that they not only expressed IgG2c, but that there were only genome sequence reads for the IgG2c gene. Likewise, strains with either A/J, 129S1, NZO, CAST, WSB, or PWK haplotypes at IgH only contained genome sequence reads for the IgG2a gene and not IgG2c. These data indicate that IgG2a and IgG2c indeed represent two distinct alleles in the mouse genome. It has been noted that there are functional differences between IgG2a and IgG2c (Petrushina et al., 2003), and these results can help investigators identify relevant mouse strains with specific IgG2 subtypes for functional follow-up to these two alleles.

B cells play a critical role in immune system development and homeostasis. Specific subsets of B cells, B1 and marginal zone B cells, differentiate to antibody secreting cells to produce natural antibodies, which are present before antigen stimulation and provide a first line of defense against infection (Holodick et al., 2017; Zhou et al., 2007; Subramaniam et al., 2010; Jayasekera et al., 2007). Natural antibodies are characterized by their broad reactivity and low affinity and are pre-existing or immediately secreted upon stimulation or a ‘light push’ (Holodick et al., 2017). These antibodies are largely thought to bridge the gap between innate and adaptive immunity and have been studied for their ability to protect against various pathogens (Panda et al., 2015; Jayasekera et al., 2007). Several aspects of natural antibody have been investigated, but it is still largely unclear how these antibodies are regulated and under what contexts they are produced (New et al., 2016; Holodick et al., 2017). Here we identify a novel genetic regulator, *Mbd1*, of pre-existing antibody, marginal zone B cell, and antibody secreting cell levels at homeostasis. Our data suggests that MBD1 inhibits marginal zone B cell differentiation to antibody secreting cells at homeostasis, thus regulating pre-existing antibody levels.

MBD1 is a known epigenetic regulator, facilitating chromatin remodeling through various protein-protein interactions, binding directly to methylated DNA, and facilitating transcriptional repression (Fujita et al., 2000; Ichimura et al., 2005). Much of what is known about MBD1’s function comes from studies of neural stem cell differentiation (Lax et al., 2017) and adipocyte differentiation (Matsumura et al., 2015). In the immune system, previous work has shown that MBD1 facilitates tissue-specific antigen expression through protein-protein interactions with AIRE to promote T cell tolerance (Waterfield et al., 2014). However, MBD1 has not been described as having a role in B cell differentiation. Chromatin modifying complexes and other epigenetic regulators are critical to cellular differentiation, as chromatin accessibility changes over differentiation states allows for proper gene expression. Recent work has highlighted the important role of epigenetic regulators in B cell activation and differentiation. For example, EZH2 (Herviou et al., 2019), LSD1 (Haines et al., 2018), and DNA methylation (Barwick et al., 2016; Barwick et al., 2018) have all been shown to regulate some aspect antibody secreting cell differentiation. Given the role of MBD1 in cellular differentiation in other tissues as well as our data presented here, it is likely that MBD1 is regulating chromatin dynamics necessary for altering gene expression profiles that promote antibody secreting cell differentiation. Thus, understanding the role of MBD1 in regulating gene expression profiles necessary for B cell differentiation will be important for defining those genes and regulatory networks that control B cell responses at homeostasis, as well as following infection and activation of adaptive immune responses. serving a comparable role.

Our study encompasses experiments using several cohorts of CC strains with varying ages and experimental designs, as well as a new *Mbd1* knockout model on the B6 background. While we find that the B6 allele is consistently associated with greater antibody levels and lower splenic B cell proportions in the CC, we were not able to recapitulate all those phenotypes in our knockout studies. For example, in the CC the B6 haplotype is not only associated with greater levels of IgG1 in the serum but also greater levels of total IgG and marginally associated with greater levels of other antibody subtypes. However, we were only able to recapitulate the impact of MBD1 on IgG1 levels, specifically, in the context of our knockout model. It is not entirely surprising that we are not able to completely recapitulate every phenotype observed in the larger initial screen. In the CC, allelic variants across many genetic backgrounds are averaged, while a knockout represents an extreme abrogation in the context of a single genetic background. In inbred B6 mice, the evolutionarily derived allele of *Mbd1* has co-evolved with protein binding partners and regulatory networks, whereas in the CC, those co-evolved networks have been broken apart and alleles are shuffled out of context. Additionally, there are 2 other founder strain haplotypes that were associated with greater antibody levels in the CC, and we do not capture their contributions in our knockout studies. Lastly, we do not map 100% of the genetic regulators of IgG1, as *Qih1* only accounts for ∼75% of the genetic regulation of IgG1 levels at homeostasis.

Although, *Qih1/Mbd1* has a large effect, there are other genes that contribute to the regulation of IgG1 and marginal zone B cell differentiation at homeostasis. None the less, we find a consistent role for *Qih1/Mbd1* on homeostatic antibody levels and various B cell subsets across our experimental populations. Specifically, we found that a functional B6 *Mbd1* allele is consistently associated with greater levels of homeostatic antibody and lower levels of various B cell subsets in the spleen. Interestingly, the association between the B6 allele and lower levels of B cell subsets was not observed in the bone marrow (**supplemental figure 1**), suggesting that overall B cell development was intact.

In summary our data provide evidence for strong genetic regulation of homeostatic immunity. We also show the utility of forward genetic screens in diverse mouse populations for identifying novel genes regulating homeostatic immunity. To our knowledge this is the first demonstration that MBD1 may act as a negative regulator of marginal zone B cell differentiation to antibody secreting cells, thereby regulating antibody levels at homeostasis. Additionally, our data further illustrates the role of epigenetic regulators in cellular differentiation and function. Future work to elucidate the specific pathways controlled by MBD1 to regulate marginal zone B cell differentiation could enhance our understanding of marginal zone B cell mediated humoral immunity at homeostasis and in response to pathogen infection.

## Methods

### Mice

#### Collaborative Cross mice

CC mice were obtained from the UNC Systems Genetics Core Facility at UNC Chapel Hill between 2013 – 2017. All experiments were approved by the UNC Chapel Hill Institutional Animal Care and Use Committee. Mice were sacrificed using isoflurane overdose and terminally bled by cardiac puncture at six to twelve weeks of age depending on the experiment. For the baseline antibody screen, experiments were conducted under biosafety level 3 (BSL3) conditions where four to six weeks old female mice were cohoused across strains and allowed to acclimate to the BSL3 for 3-4 days. Mice were then inoculated subcutaneously in the left rear footpad with phosphate-buffered saline (PBS) supplemented with 1% fetal bovine serum (FBS).

#### *Mbd1* knockout mouse

*Mbd1* knockout mice were generated at UNC Chapel Hill by the Animal Models Core via CRISPR mutagenesis. Specifically, Cas9 guide RNAs were designed with Benchling software and used to generate a 752bp deletion spanning exons 11 and 13, which resulted in protein ablation. The presence of the knockout or wildtype allele was determined by amplifying across the region using the following primers: forward primer – GCTCACTGAGTAGGGCAAGG, reverse primer – TACGGAGCACACCTTGGCA. Wildtype amplicon: 1262bp. Knockout amplicon: 510bp. We maintained these mice in our colony via 2 generations of backcrossing to C57BL/6J mice (JAX stock #000664) to remove any potential off-target mutations. The stock was thereafter maintained via het-by-het crosses.

#### ELISA

Total antibody levels were quantified by ELISA. 96-well flat-bottom high-binding plates were coated with anti-Ig antibodies (Southern Biotech) diluted in carbonate buffer for each antibody subtype measured. Serum was diluted in ELISA wash buffer (1x PBS + 0.3% Tween20) with 5% nonfat milk and added to pre-coated plates to incubate in humid storage overnight at 4°C. Plates were washed using ELISA wash buffer and incubated with HRP-conjugated anti-Ig secondary antibodies (Southern Biotech) for 2 hours in humid storage at 4°C. Plates were washed and developed in the dark for 30 minutes at room temperature with citrate substrate buffer, the reaction was stopped with sodium fluoride, and read immediately at 450nm. Standard curves for each antibody subtype (Invitrogen standards) were run with each plate for a specific subtype to determine antibody concentrations.

#### Sample preparation

Following euthanasia (as described above), blood was collected immediately into serum separator tubes to isolate serum for ELISAs. Serum was aliquoted in 1.7mL Eppendorf snap-cap tubes and stored at -80°C until analyzed by ELISA. Spleens were homogenized using frosted microscope slides and pelleted by centrifugation (1000 RPM/4°C/10 minutes). Spleen homogenates were filtered through 70um mesh, treated with ammonium chloride potassium (ACK) lysing buffer to remove red blood cells, washed, and resuspended in FACS buffer (HBSS + 1-2% FBS). Cell numbers were determined by trypan blue exclusion using a hemocytometer or Countess II automated cell counter.

#### Flow Cytometry

Cells were plated at 1-2×10^7^ per mL in FACS buffer (HBSS + 1-2% FBS) in a 96-well polypropylene round-bottom plate. Cells were centrifuged at 1000 RPM for 4 minutes at 4°C and resuspended in 100uL of fluorochrome-conjugated antibody dilution. Cells were incubated at 4°C for 45 minutes to allow for antibody staining. Following incubation, cells were washed twice with FACS buffer and resuspended in 100uL of FACS buffer.

An equal volume of 4% PFA (in PBS) was added to cells to fix, and plates were stored in the dark at 4°C until analyzed of Attune NxT flow cytometer. Data was analyzed using FlowJo software. The following antibodies were used: Live/Dead Fixable Aqua, CD3-PE (145-2C11), CD4-APC/Cy7 (GK1.5), CD8a-PerCP(53-6.7), CD11b-eF450 (M1/70), CD11c-PE/TxRd (N418), CD19-AF647 (6D5), CD45-AF700 (30-F11), CD19-APC/Cy7 (6D5), IgM-PE (ll/41), IgD-FITC(11-26c.2a), CD5-BV421 (53-7.3), CD3-PerCP (145-2C11), CD21/CD35-PE (7E9), IgM-AF594, CD23-PE/Cy7 (B3B4), CD45R-APC (RA3-6B2).

#### Data processing

Antibody concentrations were determined from a standard curve and log10 transformed to follow a normal distribution. Event counts from flow cytometry gating were used to calculate cell proportions. Using Box-Cox transformation (MASS package in R, version 3.5.1), values were independently transformed for each phenotype to follow a normal distribution. For all phenotypes used for QTL mapping, the average phenotype value was calculated for each Collaborative Cross strain and used to map.

#### Nested linear models

Correlations between identified QTL and immune cell populations and other antibody subtypes were determine by comparing the goodness of model fit of mixed effect linear models using a partial fit F-test. *Qih1* haplotype scores were determined by the founder haplotype at the *Qih1* peak marker. In both the base and full model, CC Strain is a random effect variable and *Qih1* haplotype score is a fixed effect variable. The full model tests whether including information about the haplotype at *Qih1* explains more of the phenotypic variation than the CC Strain alone.

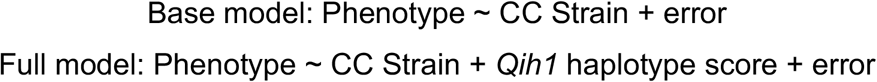

#### Heritability calculations

Heritability calculations were performed as described previously (Noll et al., 2020). Briefly, box-cox transformed phenotype values were used to fit a linear fixed-effect model. The coefficient of genetic determination was calculated as such:

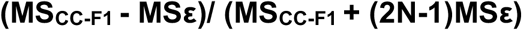

Where MSCC-F1 is the mean square of the CC-F1 and MSε is the mean square of the error using a N=3 as an average group size, as a measure of broad-sense heritability.

#### QTL mapping

QTL mapping was performed as previously described (Noll et al., 2020). Briefly, we used the DOQTL (Gatti et al., 2014) package in the R statistical environment (version 3.5.1). A multiple regression is performed at each marker, assessing the relationship between the phenotype and the haplotype probabilities for each strain. LOD scores are calculated based on the increase in statistical fit compared to a null model, considering only covariates and kinship. To calculate significance thresholds, permutation tests were used to shuffle genotypes and phenotypes without replacement. We determined the 80^th^, 90^th^, and 95^th^ percentiles after 500 permutations as cutoffs for suggestive (both p<0.2 and p<0.1) and genome-wide significant (p<0.05). QTL intervals were determined using a 1.5 LOD drop.

## Acknowledgements

This work was supported, in part, by U19AI100625 (to MTH, MTF, and FPMV), P01AI132130 (to FPMV and MTF), R21AI119933 (to MTF), HHMI Gilliam Fellowship (to BKH), and F99AG073570 (to BKH). We wish to thank the Systems Genetics Core Facility for the maintenance of the Collaborative Cross lines used in these studies. We would also like to thank the UNC Flow Cytometry Core for their help with flow cytometry data acquisition and analysis.

Studies were designed by BKH KSP ACW FPMV MTF MTH

Experiments were conducted by BKH KSP ACW CLL EAM GDS TAM PH KEN SPB

Data were analyzed by BKH ACW CLL MTF MTH

Manuscript was written primarily by BKH MTF MTH

All authors contributed to the editing of the manuscript

