## Supplemental figures and tables for "Forward genetic screen of homeostatic antibody levels in the Collaborative Cross identifies MBD1 as a novel regulator of B cell homeostasis"

### SUPPLEMENTAL FIGURES & TABLES

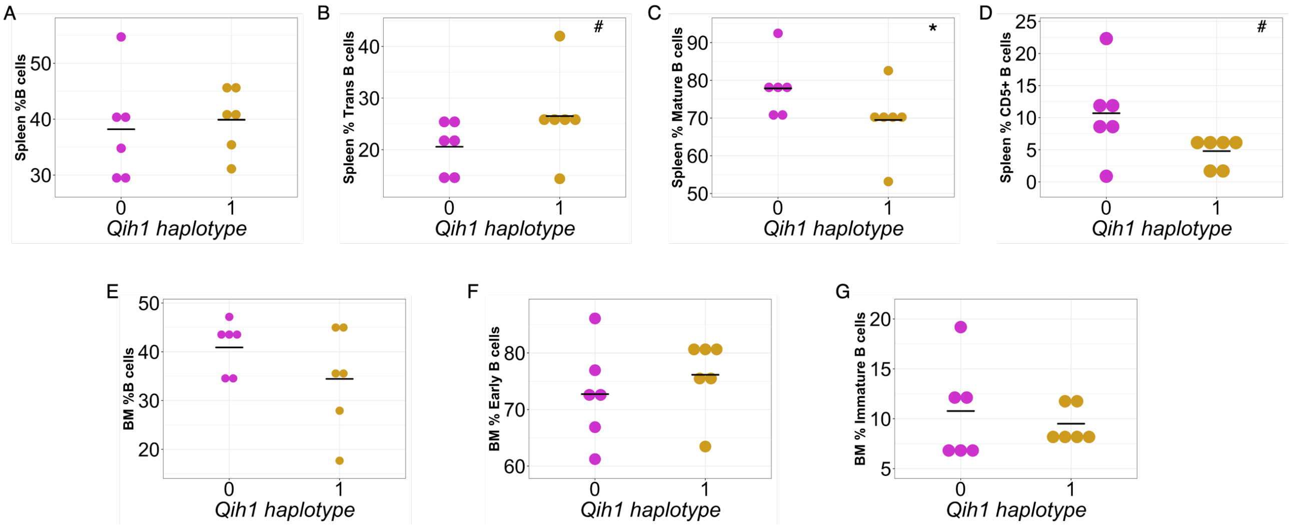

Supplemental figure 1: *Qih1* effects on B cell populations in the spleen and bone marrow. We assessed the relationship between B6, WSB, and CAST haplotypes (*Qih1* haplotype = 1) and B cell subsets in the spleen and bone marrow. Each point represents the mean value for each CC strain and the mean for each haplotype group on the x-axis is denoted by the grey crossbar. ($^{\#}p < 0.2$, $^{*}p < 0.1$)

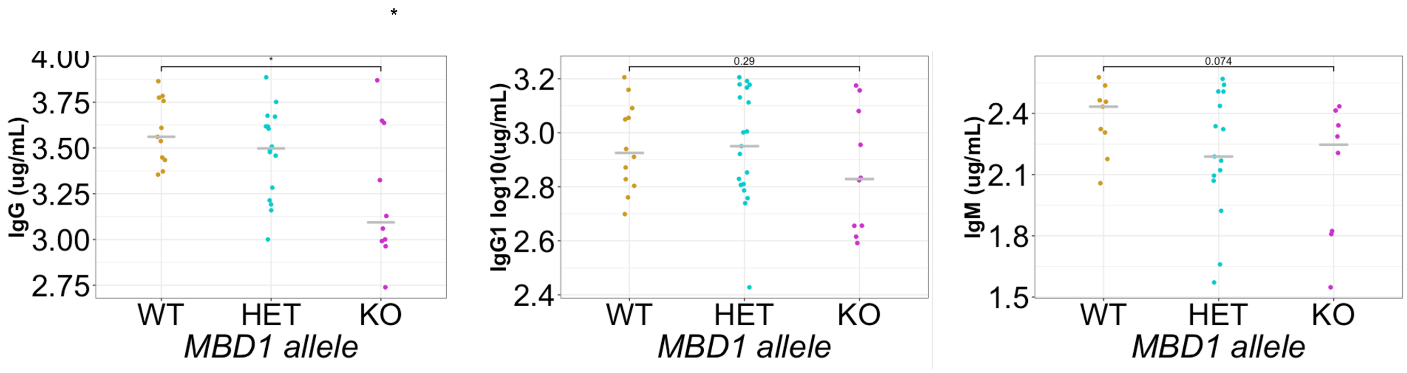

Supplemental figure 2: *Mbd1* KO mice have lower levels of antibody with increased age. We assessed IgG1, Total IgG, and IgM levels in the serum of 15-16wk old animals. Each point represents an individual animal and the median for each genotype group on the x-axis is denoted by a crossbar. ($^{*}p < 0.05$)

Supplemental Table 1: CC and inbred mouse strains used in these studies.

| Strains used in homeostatic antibody screen | Strains used in 48CC screen | Strains used in *Qih1* validation study | Strains used for MBD1 variant analysis |
|---|---|---|---|
| CC001/Unc | CC001/Unc | CC002/Unc | 129P2/OlaHsd |
| CC002/Unc | CC002/Unc | CC003/Unc | 129S1/SvImJ |
| CC003/Unc | CC003/Unc | CC004/TauUnc | 129S5SvEvBrd |
| CC004/TauUnc | CC004/TauUnc | CC008/GeniUnc | A/J |
| CC006/TauUnc | CC005 | CC013/GeniUnc | AKR/J |
| CC007/Unc | CC007/Unc | CC031/GeniUnc | BALB/cJ |
| CC008/GeniUnc | CC008/GeniUnc | CC032/GeniUnc | BTBR/T/Itpr3tf/J |
| CC009/Unc | CC011/Unc | CC037/TauUnc | BUB/BnJ |
| CC010/GeniUnc | CC013/GeniUnc | CC045/GeniUnc | C3H/HeH |
| CC011/Unc | CC015/Unc | CC058/Unc | C3H/HeJ |
| CC012/GeniUnc | CC016/GeniUnc | CC062/Unc | C57BL/10J |
| CC013/GeniUnc | CC017/Unc | CC074/Unc | C57BL/6NJ |
| CC015/Unc | CC019/TauUnc | | C57BR/cdJ |
| CC016/GeniUnc | CC021/Unc | | C57L/J |
| CC017/Unc | CC023/GeniUnc | | C58/J |
| CC018/Unc | CC024/GeniUnc | | CAST/EiJ |
| CC019/TauUnc | CC025/GeniUnc | | CBA/J |
| CC021/Unc | CC026/GeniUnc | | DBA/1J |
| CC022/GeniUnc | CC027/GeniUnc | | DBA/2J |
| CC023/GeniUnc | CC029/Unc | | FVB/NJ |
| CC024/GeniUnc | CC030/GeniUnc | | I/LnJ |
| CC025/GeniUnc | CC031/GeniUnc | | KK/HiJ |
| CC026/GeniUnc | CC032/GeniUnc | | LEWES/EiJ |
| CC027/GeniUnc | CC033/GeniUnc | | LP/J |
| CC028/GeniUnc | CC036/Unc | | MOLF/EiJ |
| CC029/Unc | CC037/TauUnc | | NOD/ShiLtJ |
| CC030/GeniUnc | CC038/GeniUnc | | NZB/B1NJ |
| CC031/GeniUnc | CC039/Unc | | NZO/HILtJ |
| CC032/GeniUnc | CC040/TauUnc | | NZW/LacJ |
| CC033/GeniUnc | CC041/TauUnc | | PWK/PhJ |
| CC035/Unc | CC042/TauUnc | | RF/J |
| CC036/Unc | CC043/GeniUnc | | SEA/GnJ |
| CC037/TauUnc | CC044/Unc | | SPRET/EiJ |
| CC038/GeniUnc | CC045 | | ST/bJ |
| CC039/Unc | CC046/Unc | | WSB/EiJ |
| CC040/TauUnc | CC049/TauUnc | | ZALENDE/EiJ |
| CC041/TauUnc | CC055/TauUnc | | |
| CC042/GeniUnc | CC057/Unc | | |

| | |
|---|---|
| CC043/GeniUnc | CC058/Unc |
| CC044/Unc | CC059/TauUnc |
| CC046/Unc | CC060/Unc |
| CC049/TauUnc | CC062/Unc |
| CC051/TauUnc | CC071/TauUnc |
| CC055/TauUnc | CC072/TauUnc |
| CC056/GeniUnc | CC078/Unc |
| CC057/Unc | CC080/Unc |
| CC058/Unc | CC081/Unc |
| CC059/TauUnc | CC082/Unc |
| CC060/Unc | |
| CC061/GeniUnc | |
| CC062/Unc | |
| CC065/Unc | |
| CC068/TauUnc | |
| CC070/TauUnc | |
| CC071/TauUnc | |
| CC072/TauUnc | |
| CC074/Unc | |
| CC075/Unc | |

Supplemental Table 2: Phenotype distribution and heritability of 48CC screen phenotypes.

| Phenotype | Phenotypic range | Median Phenotype Value | Heritability Estimate |
|---|---|---|---|
| CD3+ T cells | 0.113 – 0.678 | 0.342 | 0.493 |
| CD4+ T cells (% of CD3+ T) | 0.339 – 0.782 | 0.606 | 0.761 |
| CD8+ T cells (% of CD3+ T) | 0.121 – 0.489 | 0.269 | 0.706 |
| CD19+ B cells | 0.246 – 0.622 | 0.441 | 0.243 |
| CD11b+ cells | 0.010 – 0.160 | 0.038 | 0.308 |
| CD11c+ cells | 0.007 – 0.968 | 0.022 | 0.428 |
